# Sexual Dimorphism in Jump Kinematics and Choreography in Peacock Spiders

**DOI:** 10.1101/2024.08.09.607331

**Authors:** Ajay Narendra, Anna Seibel, Fiorella Ramirez-Esquivel, Pranav Joshi, Donald James McLean, Luis Robledo-Ospina, Dinesh Rao

## Abstract

Jumping requires a rapid release of energy to propel an animal. Terrestrial animals achieve this by relying on the power generated by muscles, or by storing and rapidly releasing elastic energy. Jumping spiders rely on hydraulic pressure and muscular action to propel their jump. Though males and females of jumping spiders vary in size, sex-specific differences in jumping have never been studied. We investigated sexual dimorphism in the jump kinematics of an Australian Peacock spider, *Maratus splendens*. We recorded locomotory jumps in males and females using high-speed videography (5000 frames per second). We determined the centre of mass of the animals using µCT and tracked its displacement during a jump. We found that although females weighed more than twice as much as males, both had similar accelerations and take-off velocities. Males had shorter jump take-off duration, steeper take-off angle and experienced higher g-force compared to the females. We examine the jump choreography of male and female spiders and explore the factors behind the differences in their jump kinematics.

## Introduction

To execute a jump, animals require a rapid burst of energy to launch themselves from a substrate, during which they coordinate various body parts to move synchronously within a brief period. Jumping is an energetically expensive locomotory strategy [1, 2] that rapidly moves an animal either towards or away from a target. Many taxa have developed distinct strategies for jumping or leaping, including mammals, marsupials, amphibians and arthropods [3–7]. Animal jumps are classified as muscle-actuated or spring-actuated systems based on how they are powered. Muscle-actuated jumps are seen in ants [8], lacewings [9], and bush crickets [10], where animals rely on the power generated by the contraction of leg muscles (e.g., muscles in the coxae, trochanter, and tibia). The volume of the muscle and its physiological properties limits the jump performance [8, 11]. Spring-actuated systems, also referred to as catapult systems or power-amplification systems, as seen in crickets, click beetles, true bugs and fleas, bypass muscular limitations and output more power by relying on internal structures to store energy which is then rapidly released to trigger the jump [12, 13].

Jumping spiders of the family Salticidae, are exceptional jumpers as their name suggests. They leap to cross gaps, navigate detours, capture prey, and evade predators [12–17]. Salticids jump to targets at different elevations and often cross gaps 10x-15x times their body length [17]. Spider jumps are neither muscle nor spring actuated, instead they use an unusual strategy referred to as the semi-hydraulic system [15, 18–20]. Spider legs have flexor muscles, which are used for postural movements, but lack the extensor muscles required to extend their legs and power jumps. Instead, the legs are extended by increasing the haemolymph pressure in the legs. This pressure is generated by the contraction of muscles in the cephalothorax, which pumps haemolymph into the leg. Since our understanding of how spiders control the semi-hydraulic system is relatively limited, there are no clear predictions on how jump kinematics in spiders scale with size.

Currently, jumps have been described only in three genera of Salticidae: *Attulus, Habronattus and Phidippus* [15, 16, 21, 22]. In most Salticids, males are smaller compared to females who must build up nutritional reserves to produce eggs. While behavioural tasks that require jumping are similar between the two sexes, there is one distinct difference: males engage in rapid escapes following mating to avoid female attacks and to minimise competition from other males.

Here, we investigated the jump kinematics and choreography of males and females of an Australian Peacock spider, *Maratus splendens* (Fig. 1A, 1B). Peacock spiders are native to Australia, with males renowned for their vibrant iridescent colours and elaborate courtship displays [23, 24]. During courtship, males of *Maratus* species spread open their brightly coloured opisthosomal flaps, which are usually folded around the abdomen, and oscillate around a female while waving the ornamented 3^rd^ pair of legs [23, 25, 26]. To the best of our knowledge, sex-specific differences in Salticid jumps have not been investigated. Using high-speed videography, we studied locomotory jumps and determined the jump kinematics and choreography in male and female spiders.

**Figure 1.**
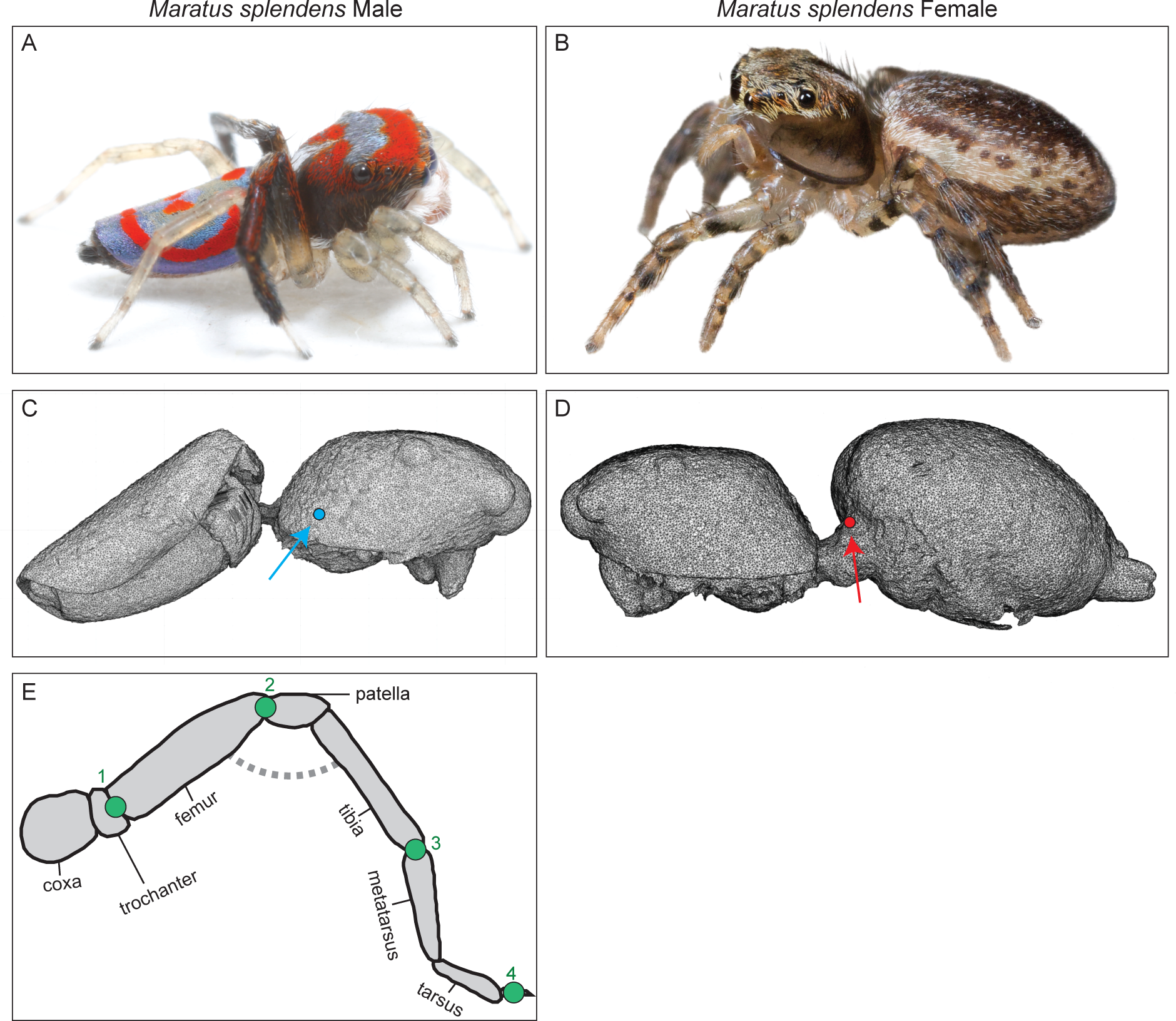
Male and female of the Australian Splendid Peacock spider, *Maratus splendens*. (A, B) Colour photographs of the male and female spider. (C, D) 3D render from µCT scans of male and female spiders. The centre of mass without legs is illustrated by a circle in male (red) and female (blue) spiders. (E) Schematic of the spider’s leg, illustrating the four joints that were tracked: (1) trochanter-femur, (2) femur-patella, (3) tibia-metatarsus, and (4) tarsal claw. We determined Effective Leg Length on every frame as the distance between trochanter to tarsal claw by sum of length of each segment. The flex in the joint on each frame was determined as the angle between trochanter-femur and femur-patella segments. Photo credits: Male: Ajay Narendra, Female: Jürgen Otto

## Methods

We collected adult males and females of the Splendid Peacock spider, *Maratus splendens* [27] at the Lake Paramatta Reserve, Sydney, New South Wales, Australia. Spiders were individually housed in containers and provided water and food (*Drosophila melanogaster*). Experiments were carried out between October-December 2023 at the Wallumattagal campus, Macquarie University.

### Experimental setup and filming

Experiments were carried out in the laboratory with flicker free LED lighting, and constant temperature conditions of 24°C. We fixed a landing and take-off platform to an optical bench which was placed on a vibration isolation table. The platforms were 3D printed with black PLA using Ultimaker S5. The take-off platform (4 x 0.5 x 0.1 cm, L x W x H) was horizontal to the ground to which a spider walked on their own from a collection jar. We placed a vertical landing platform (0.1 x 0.5 x 4 cm, L x W x H) 4 cm away from the take-off platform. Individual spiders jumped from the take-off to the landing platform of their own accord (Fig. 2). Thus, jumps were not elicited but allowed to happen naturally. Some spiders jumped immediately, whereas others took a few minutes to jump towards the landing platform. We removed all other visual targets in the vicinity, which ensured that spiders carried out locomotory jumps to the landing platform. For each individual, we recorded three consecutive jumps. For one female spider we only recorded two jumps, as this animal was reluctant to jump to the target on its third attempt. Animals were weighed immediately after they carried out the jumps (Sartorius Entris ii Advanced line Balance, Göttingen, Germany, 0.1 mg repeatability) and subsequently released back to their natural habitat.

**Figure 2.**
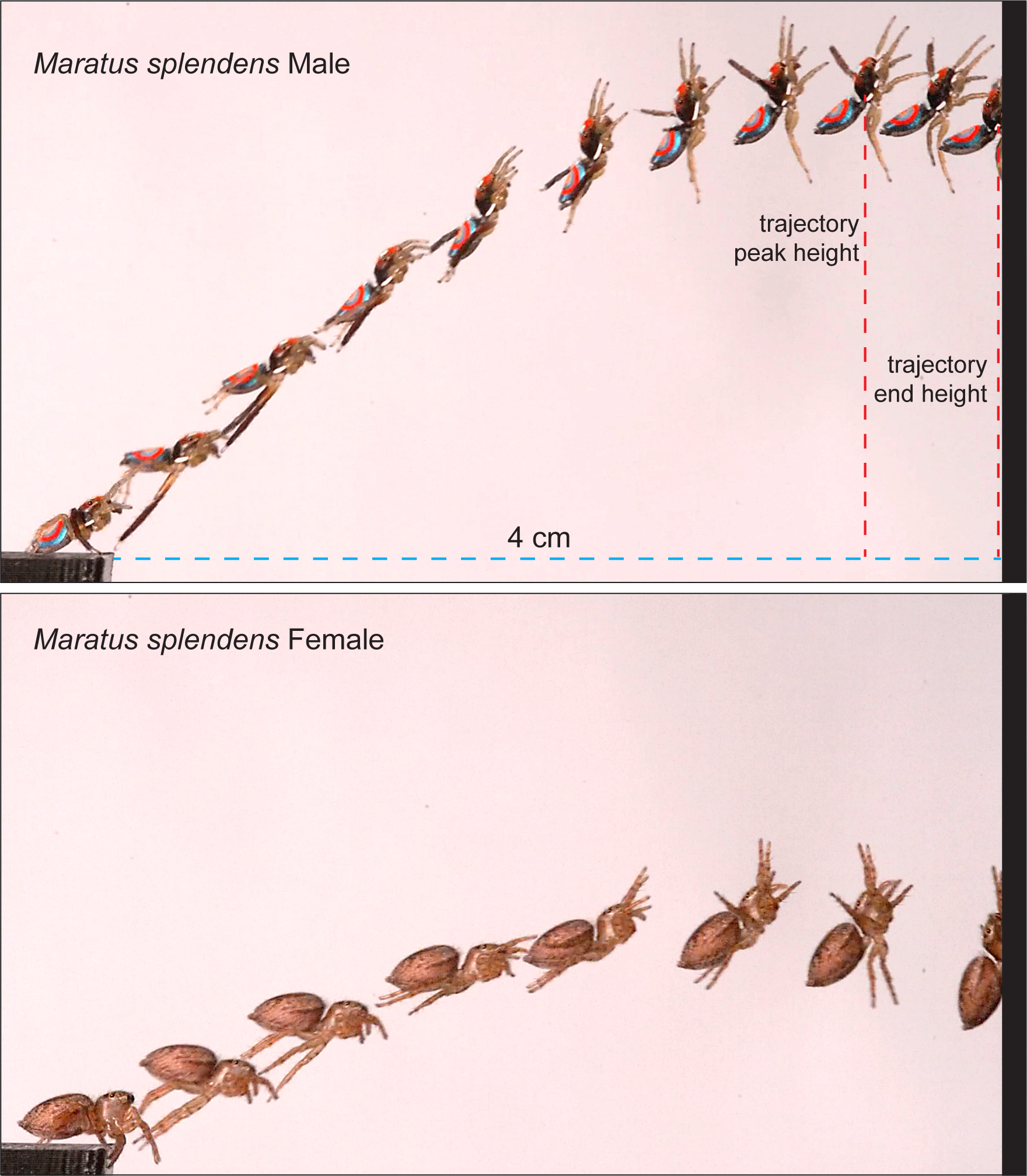
Experimental setup and jump sequence in the male and female Australian Splendid Peacock spider, *Maratus splendens*. Individual spiders jumped on their own accord from the take-off platform on the left to cross a 4 cm gap to reach a vertical landing platform. Spiders were filmed in profile at 5000fps. Time lapse series to illustrate the jump is shown for an example male (top) and female spider. The images chosen are for clarity and are not spaced at regular timepoints. Two vertical dashed lines illustrate the trajectory peak height and trajectory end height.

We filmed the jumps with a 105mm Sigma lens attached to a high-speed camera (T1340, Phantom, Adept Turnkey Pty Ltd, Australia) at 5000fps (2048 x 1024 pixels). The spider and filming area was illuminated with Godox SL 300III light source. We filmed either the entire jump trajectory (male: n=4 spiders; 12 jumps; female: n=7 spiders; 20 jumps), or magnified views of take-off jumps to track leg joints accurately (male: n=6 spiders; 18 jumps; female: n=5 spiders; 15 jumps).

Raw footage was converted from cine to uncompressed mp4 files and trimmed in Videoloupe (Corduroy Code Inc. British Columbia, Canada). Trimmed files were imported to DLTdv8 [28] in Matlab 2023b (Mathworks, Nattick, Massachusetts) where we carried out a manual frame-by-frame analysis to track different body structures and extract x,y, coordinates. In both full-jumps and take-off only jumps we tracked the centre of mass (CoM; see below). From these coordinates, we constructed the jump trajectory. In full jumps, we identified the maximum height attained during jump and the height above ground when animals reached the landing platform. We used the displacement of the CoM to determine jump kinematics, which included:

i. Take-off duration (ms) = duration between the first instance when acceleration increased from zero to the first instance when all legs were off the ground
ii. Velocity (m/sec), *v = d/t,* where d=distance, t=time
iii. Acceleration (m/sec^2^), *a = Δv/Δt*, where Δv = change in velocity, Δt = change in time
iv. Kinetic Energy (µJ), *k.e. = ½ mv^2^*, where m=mass, v=velocity
v. Jump Force (mN), *F = m x a* (mN)
vi. Jump Power (mW), *P = k.e. / take-off duration*
vii. G-force, *g = a / 9.81*
viii. take-off angle (degrees): the angle of CoM between ‘the first frame prior to when the abdomen moves’ and ‘the first frame when all legs are off the ground’ relative to the plane of the substrate.

### Contribution of different legs for jumping

To identify the contribution of different legs in a jump, we tracked four leg joints in each leg in one randomly selected jump of all males (n=6) and female spiders (n=5) from the high magnification footage. We tracked the following joints: trochanter-femur, femur-patella, tibia-metatarsus, and tarsal claw (Fig. 1E). Over the course of the jump, we determined the extent to which the leg extends using two methods, both adapted from Brandt et al [15]. In the first, we calculated the Effective Leg Length (ELL), which was a fraction of absolute distance between trochanter and tarsal claw/sum of length of all segments. Values range between 0 and 1, where values close to 1 indicate a fully extended leg. In the second, we determined the flex, i.e., the angle between trochanter-femur segment and femur-patella segment. Lastly, we also determined the time difference between when animals reached peak acceleration and when all legs had left the ground.

### Identifying the centre of mass

Specimens were fixed in 4% paraformaldehyde for 4 hours and then rinsed in 0.1M phosphate buffer (3 x 10 minutes). Specimens were stained in 2% Lugol’s solution for 48 hours followed by washes in a buffer (3 x 10 minutes). Samples were individually placed in small Eppendorf tubes filled with 70% ethanol for imaging for µCT. We imaged the samples with a Bruker SkyScan1272 microCT scanner using a voltage of 45kV, a current of 200μA, and a 0.5mm aluminium filter. Projection images were reconstructed using the software SkyScan’s NRecon v.2.2.0.6 software (Bruker, Kontich, Belgium). Scans achieved a voxel resolution of between 1.5 – 5.0 µm.

Image segmentation and mesh generation was carried out using ITK-snap 4.0.0 [29]. Mesh clean-up and repair were carried out using MeshLab v2022.02 [30]. Centre of mass was determined using AutoDesk Fusion 360 2.0.17721. The legs were removed for the analysis and both the cephalothorax and abdomen were assumed to have the same density.

#### Statistical Analyses

We used circular statistics, the Watson-Williams test in Oriana (Kovach Computing Services), to determine whether the mean take-off angles differed between the male and female spiders. We used linear mixed-effects models to examine the relationship between morphometric measures, the sex of the spiders, and kinematic variables. The modelling was carried out in R using the “lme4” and “lmerTest” packages. The initial analyses involved fitting a full model with random slopes and intercepts attributable to the sex of the spiders, along with a fixed effect for body size. The random effects structure was determined based on Akaike’s Information Criterion (AIC) by comparing the fits of the two models, one with a random slope, and intercept attributable to the sex of the spider and one with just sex as the random intercept. After determining the optimal random effects’ structure, the significance of the fixed effect for body size was assessed using the F test with Satterthwaite correction as implemented in the “lmerTest” package.

## Results

### Morphology

Males of *M. splendens* were distinctly lighter by weight compared to their female counterparts (male: 4.94 ± 1.86 mg, n=10; female 10.40 ± 2.05 mg; mean ± s.d.; Mann Whitney, U=3.0, p<0.001, Table 1). The centre of mass (CoM) of both sexes was determined from a µCT scan volume data in the absence of legs and assuming constant density. The CoM in the male was located on the cephalothorax, just above the fourth coxae (Fig. 1C). The CoM in the female was located on the anterior-most region of the abdomen (Fig. 1D).

**Table 1.** Jump kinematics in Salticidae. Values shown are average values for each criterion. For *Habronattus* we only used data with jump gaps of 4 cm and 5 cm (15). For *Phidippus regius* we only used data with 3, 4.5 and 6 cm gaps which were in the same horizontal plane (16). Data for *Attulus pubescens* were derived from Parry and Brown (22) and *Phidippus princeps* from Hill (21).

| Species<br>Sex<br>N<br>Gap size | Mass<br>(mg) | Acceler-<br>ation<br>(m/s<sup>2</sup>) | Velocity<br>(m/s) | Take-off<br>time<br>(ms) | Take-off<br>angle<br>(deg) | KE<br>(μJ) | Jump<br>force<br>(mN) | Jump<br>power<br>(mW) | G<br>force | Inter-<br>frame<br>interval<br>(ms) |
| --- | --- | --- | --- | --- | --- | --- | --- | --- | --- | --- |
| *Maratus splendens*<br>(male)<br>N=10<br>Gap: 4 cm | 4.9 | 127.8 | 0.84 | 20.8 | 25.08 | 1.7 | 0.63 | 0.09 | 13.03 | 0.2 |
| *Maratus splendens*<br>(female)<br>N=12<br>Gap: 4 cm | 10.3 | 122.7 | 0.81 | 24.6 | 20.89 | 3.4 | 1.25 | 0.15 | 12.5 | 0.2 |
| *Attulus pubescens*<br>(male)<br>N=2<br>Gap: 5 cm | 10 | 51.3 | 0.67 | 13.06 | 12 | 2.24 | 0.51 | 0.17 | 5.23 | 11.9 |
| *Habronattus conjunctus*<br>(mixed)<br>N=9<br>Gap: 4-5 cm | 14.5 | 28.3 | 0.48 | 17.65 | 18.7 | 1.76 | 0.40 | 0.10 | 3.18 | 1 |
| *Phidippus princeps*<br>(female)<br>N=1<br>Gap: 6 cm | 150 | 51.4 | 0.83 | 16.15 | 22 | 51.6 | 7.71 | 3.20 | 5.2 | 15 |
| *Phidippus regius*<br>(female)<br>N=1<br>Gap: 3-6 cm | 150 | 36.07 | 0.79 | 22.19 | 19.67 | 48.36 | 5.41 | 2.24 | 3.68 | 0.31 |

### Overview of jump kinematics

All male and female spiders jumped to cross a gap of 4 cm to reach a vertical landing platform (Fig. 3). We studied jump kinematics of 10 males (30 jumps) and 12 female spiders (35 jumps) and on every occasion, all animals successfully reached the landing platform.

**Figure 3.**
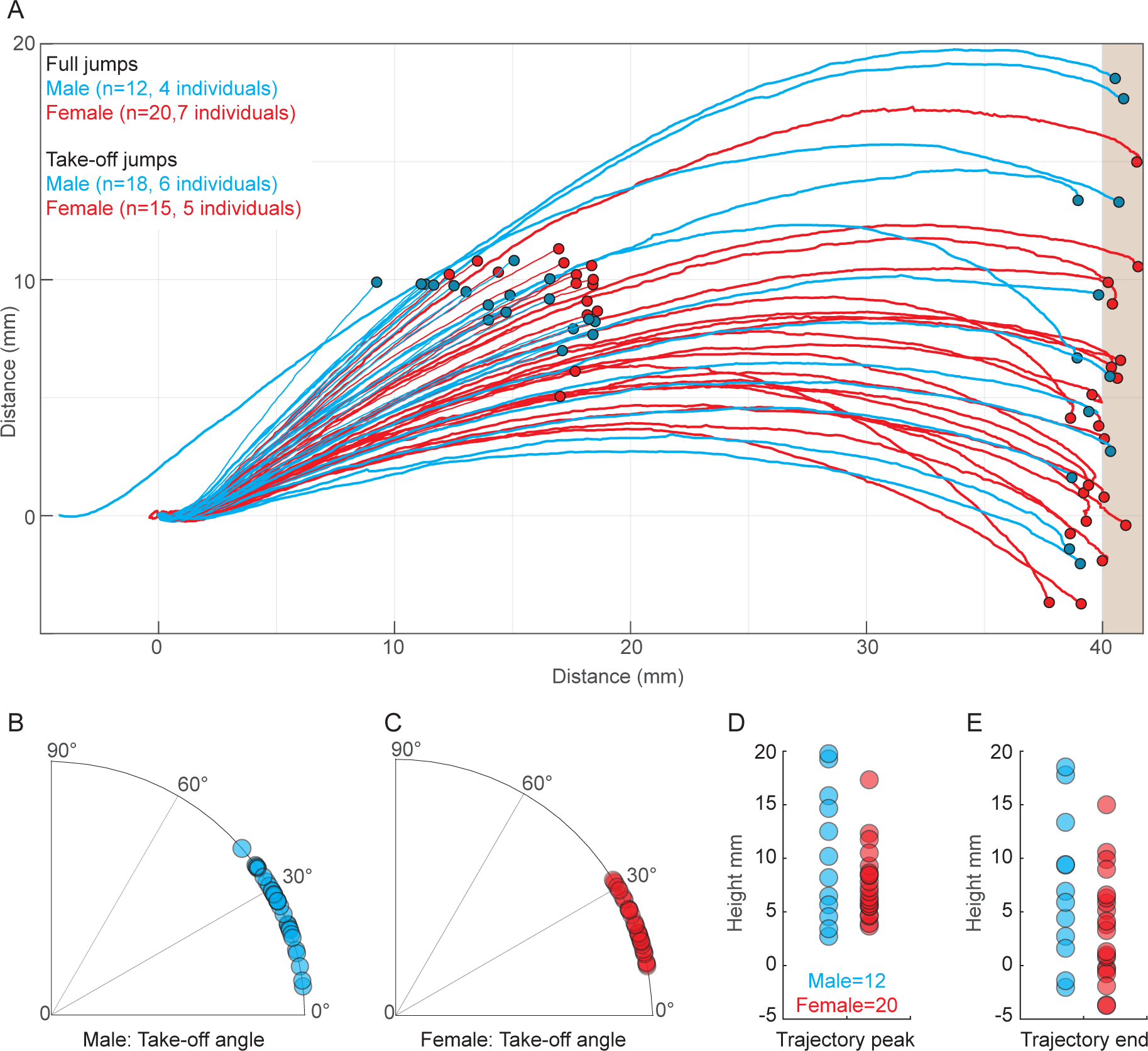
Jump trajectories and characteristics in the male and female Australian Splendid Peacock spider, *Maratus splendens*. (A) Jump trajectories start at 0,0. Spiders jumped to a vertical target placed 40 mm away. Trajectories shows are for *Full jumps*: complete trajectories from take-off to landing and *Take-off jumps*: trajectories from take-off to up to 15 cm from origin. The Centre of Mass was tracked in every frame to construct the trajectory. Endpoint of each trajectory is shown by a small circle. One male (blue line) began its jump before reaching the end of the edge of the take-off platform and hence does not begin at 0,0. (B, C) Distribution of take-off angles in males (red) and females (blue). Each circle represents take off angle of one jump. Trajectory height at the (D) peak and (E) end of the jump. For panels D and E, we only used data from full trajectories.

The smallest individual filmed was a 2.0 mg male that had a take-off duration of 22 ms, peak acceleration of 141.80 m/s^2^, and maximum velocity of 14.45 m/s. In comparison the heaviest individual filmed was a 13.2 mg female that had a take-off duration of 14 ms, peak acceleration of 122.33 m/s^2^ and a maximum velocity of 12.47 m/s. Between both sexes, the highest and lowest acceleration and velocities were seen in female spiders (acceleration: 155.22m/s^2^ and 90.06 m/s^2^; velocity: 15.82 m/s and 9.81 m/s).

### Sex and size effects on jump kinematics

Overall, males and females exhibited little variation in the maximum velocity and peak acceleration achieved during the jump and thus sex was not a significant factor (Fig. 4, Table 2). Body size, however, had a significant effect on both acceleration and velocity (Table 2). Body mass and sex had significant effect on all other kinematic variables: take-off duration, kinetic energy, jump force, jump power, g-force (Fig. 4; Tables 1, 2). Males had a significantly higher take-off angle compared to the females (males = 25.08 ± 0.9°, n=30; female= 20.89 ± 0.9°, n=35, mean ± s.d.; Watson-William test F=5.011, P=0.029; Figs. 3B, 3C, 4; Table 1). From full jump trajectories we measured the maximum height of the trajectory and trajectory height when spiders reached the landing platform (Fig. 2) and found no differences between sexes (Mann Whitney: maximum trajectory height: U=98.0, P=0.408; trajectory end height: U=98.0, P=0.146; Fig. 3D, 3E).

**Figure 4.**
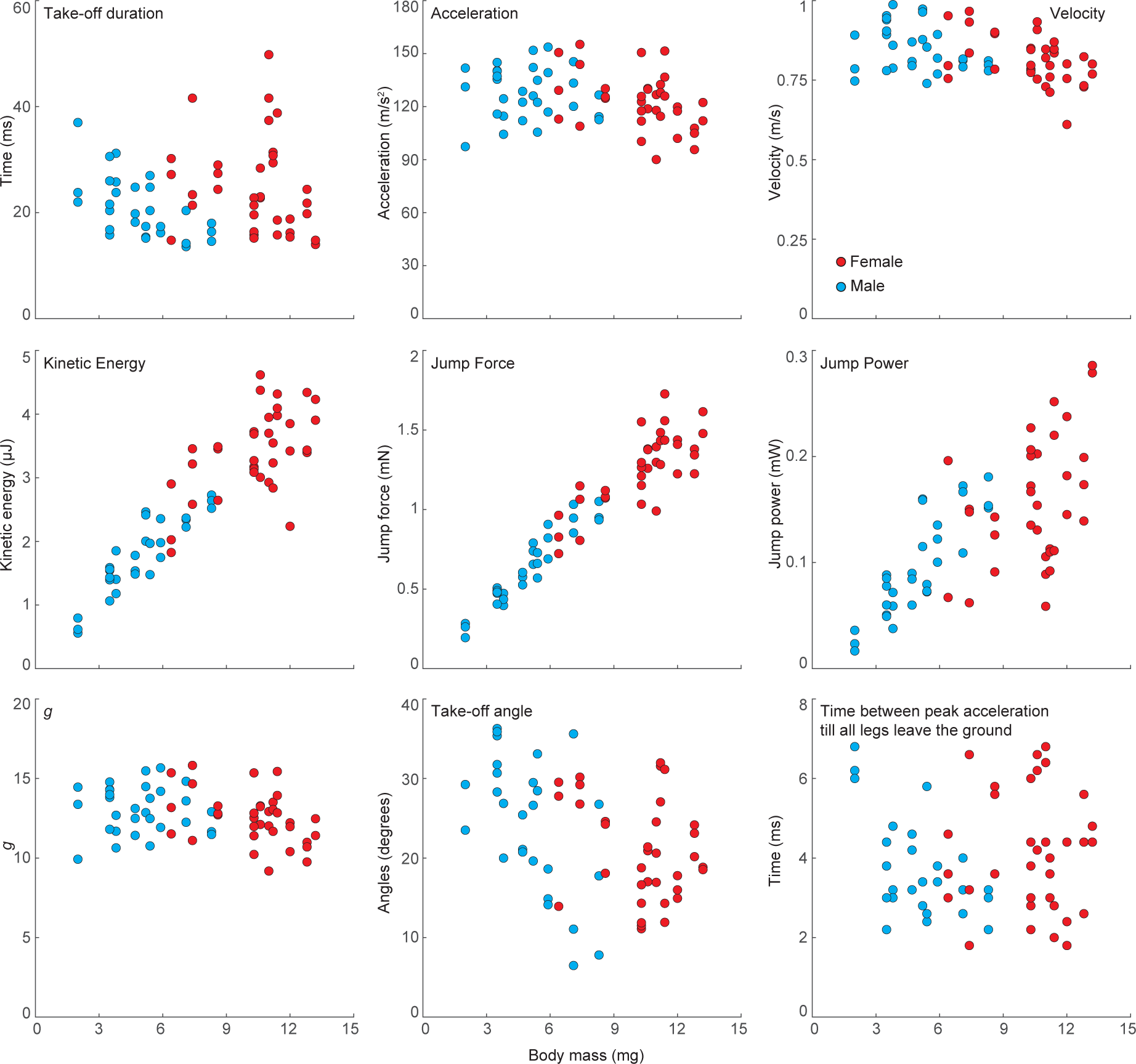
Relationship between jump kinematics and body mass in the male and female Australian Splendid Peacock spider, *Maratus splendens*. Scatter plots show relationship of body mass with take-off duration (ms), peak acceleration (m/s^2^), maximum velocity (m/s), kinetic energy, jump force, jump power, *g*, take-off angle and time duration between peak acceleration till all legs (Leg III) left the ground.

**Table 2.** Statistical outputs of the linear mixed-effects model.

| Parameters | Random Effects |  |  | Fixed Effects |  |  |  |  |  | Type III ANOVA |  |  |  |
| --- | --- | --- | --- | --- | --- | --- | --- | --- | --- | --- | --- | --- | --- |
|  | Groups | Variance | Std. dev |  | Estimate | Std. Error | df | T value | P value | Num DF | Den DF | F value | P value |
| Velocity | sex (intercept) | 0 | 0 | Intercept | -0.0651 | 0.0433 | 63 | -1.504 | p > 0.05 | 1 | 63 | 8.824 | p < 0.01 |
|  | Residual | 0.0075 | 0.0864 | Body | -0.0639 | 0.0215 | 63 | -2.971 | p < 0.01 |  |  |  |  |
| Acceleration | Groups | Variance | Std. dev |  | Estimate | Std. Error | df | T value | P value | Num DF | Den DF | F value | P value |
|  | sex (intercept) | 0.0004 | 0.0211 | Intercept | 4.8224 | 0.0214 | 0.9938 | 225.5 | p < 0.01 | 1 | 63 | 3.0908 | p > 0.05 |
|  | Residual | 0.0152 | 0.1232 |  |  |  |  |  |  |  |  |  |  |
| Jumping Force | Groups | Variance | Std. dev |  | Estimate | Std. Error | df | T value | P value | Num DF | Den DF | F value | P value |
|  | sex (intercept) | 0 | 0 | Intercept | -1.9819 | 0.0612 | 63 | -32.37 | p < 0.001 | 1 | 63 | 970.49 | p < 0.001 |
|  | Residual | 0.0149 | 0.1221 | Body | 0.9465 | 0.0304 | 63 | 31.15 | p < 0.001 |  |  |  |  |
| Kinetic Energy | Groups | Variance | Std. dev |  | Estimate | Std. Error | df | T value | P value | Num DF | Den DF | F value | P value |
|  | sex (intercept) | 0 | 0 | Intercept | -0.8234 | 0.0866 | 63 | -9.504 | p < 0.001 | 1 | 63 | 411.58 | p < 0.001 |
|  | Residual | 0.0299 | 0.1728 | Body | 0.8722 | 0.043 | 63 | 20.287 | p < 0.001 |  |  |  |  |
| Jumping Power | Groups | Variance | Std. dev |  | Estimate | Std. Error | df | T value | P value | Num DF | Den DF | F value | P value |
|  | sex (intercept) | 0.0726 | 0.2695 | Intercept | -4.497 | 0.325 | 5.014 | -13.838 | p < 0.001 | 1 | 49.635 | 76.677 | p < 0.001 |
|  | Residual | 0.1194 | 0.3455 | Body | 1.183 | 0.135 | 49.635 | 8.757 | p < 0.001 |  |  |  |  |
| Take-off Duration | Groups | Variance | Std. dev |  | Estimate | Std. Error | df | T value | P value | Num DF | Den DF | F value | P value |
|  | sex (intercept) | 0.0895 | 0.2993 | Intercept | 3.7506 | 0.3048 | 3.3892 | 12.306 | p < 0.001 | 1 | 59.762 | 9.689 | p < 0.01 |
|  | Residual | 0.0802 | 0.2832 | Body | -0.3504 | 0.1126 | 59.7617 | -3.113 | p < 0.01 |  |  |  |  |
| g | Groups | Variance | Std. dev |  | Estimate | Std. Error | df | T value | P value | Num DF | Den DF | F value | P value |
|  | sex (intercept) | 0.0004 | 0.0211 | Intercept | 2.5389 | 0.0214 | 0.9938 | 118.7 | p < 0.01 | 1 | 63 | 3.0908 | p > 0.05 |
|  | Residual | 0.0152 | 0.1232 |  |  |  |  |  |  |  |  |  |  |

### Leg kinematics

The sequence of leg and body movements between the male and female spiders were similar. Prior to jumping, spiders of both sexes raised and extended their first two pairs of legs (Figs. 2, 5; Movie S1). Spiders waved the first two pairs of legs for between 0.4 to 60 ms, and gradually lowered their abdomen. The tip of the abdomen touched the take-off platform briefly (∼0.2 ms, equivalent to interframe interval), presumably to attach a drag line thread. Legs IV left the take-off platform first, followed by Legs III (Fig. 5; Movie S1). In females, Legs III left the platform 3.5 ms (s.d = 0.4 ms, n=35) after Legs IV, whereas in males it was slightly delayed (3.8 ± 1.8 ms, mean ± s.d. n=30; Mann-Whitney U=216, P<0.001; Fig. 5; Movie S1).

**Figure 5.**
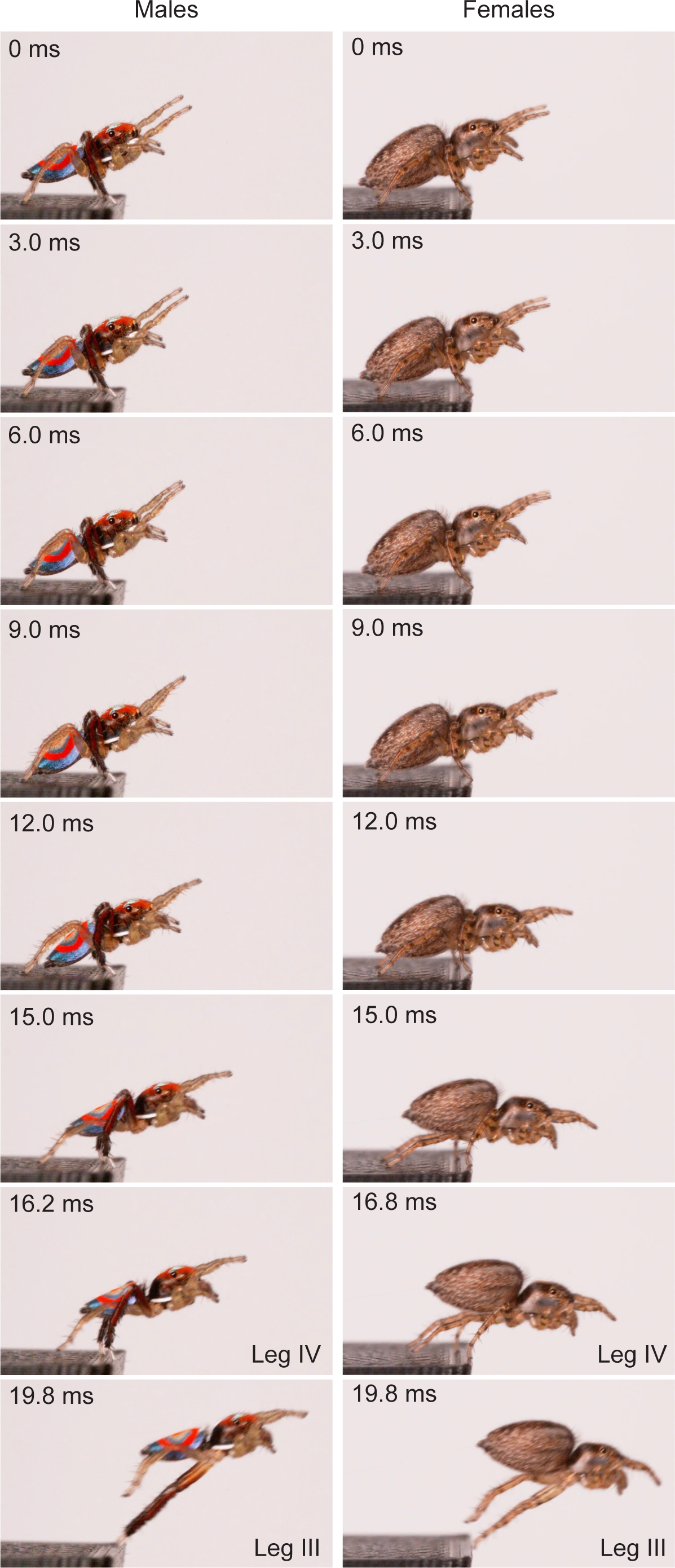
An example of a jump sequence in a male and female Australian Splendid Peacock spider, *Maratus splendens*. 0 ms: Pose held in the last frame before movement where Legs I and II raised and extended; between 3-9 ms: abdomen tip gradually moves down to touch the ground; 12 ms: Leg IV begins to extend, and CoM position moves forward; 15 ms: Leg IV is mostly extended, Leg III joint is acute, CoM position raises up. First instance when Legs IV leaves the platform (male: 16.2 ms; female: 16.8 ms) and when Legs III leave the platform (19.8 ms).

We determined the duration between peak acceleration and when all legs had left the take-off platform. In both sexes, Leg IV left the platform close to when peak acceleration occurred (females: 0.4 ± 0.2 ms, n=35 after peak acceleration; males: 0.3 ± 0.6 ms, n=30 before peak acceleration). In almost all jumps, Legs IV left the platform before peak acceleration, with the exception being a jump in one male (out of 30) and one female (out of 35). In both sexes, Leg III left the platform after peak acceleration (Fig. 5; Movie S1; males: 3.4 ± 1.4 ms, n=30 after peak acceleration; females: 4.0 ± 1.5 ms, n=35 after peak acceleration;).

In a smaller sample size (6 males; 5 females, 1 jump each) we determined the extension and the timing of leg movement by calculating the ELL and angle of the leg joint (trochanter-femur and metatarsus-tibia) relative to acceleration (Fig. 6). The ELL of Leg IV in males and females gradually increased (∼6 ms in males; ∼8 ms in females) and was fully extended before animals reached peak acceleration (with one exception in a male and female, Fig. 6). In both sexes, maximum ELL and the least acute angle in Leg III occurred after peak acceleration (Fig. 6).

**Figure 6.**
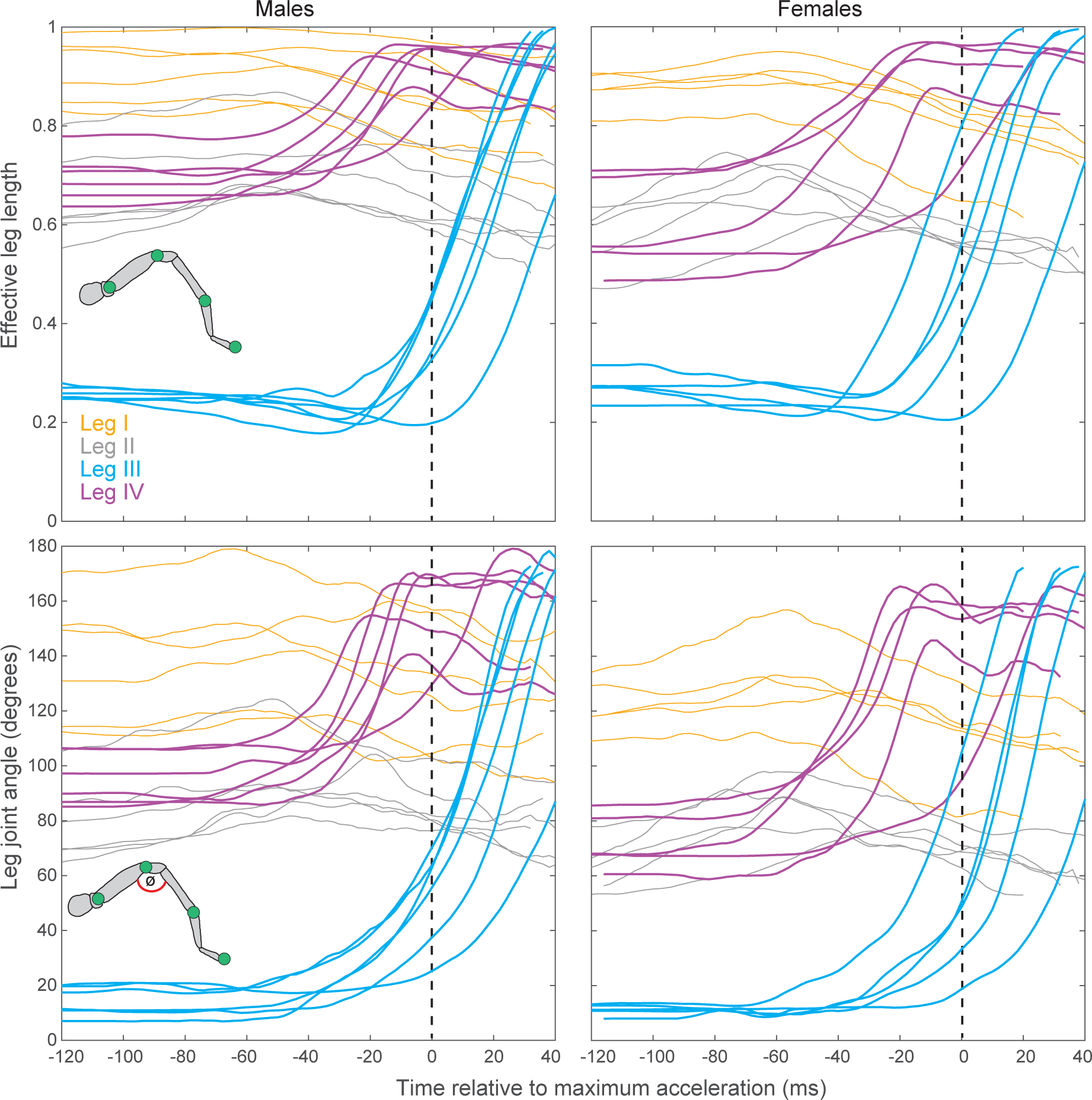
Choreography of leg movements relative to peak acceleration during a jump in the male and female Australian Splendid Peacock spider, *Maratus splendens*. Joints of four legs were tracked in 6 male and 5 female spiders. Each colour represents a leg. Inset shows schematic of spider leg with the tracked leg joints (blue circle) and the angle between trochanter-femur and femur-patella segments (ø). Top row: Effective leg length is the distance between trochanter to tarsal claw by sum of length of each segment. Values close to 1 indicate the leg is fully extended. Bottom row: The flex in the joint was determined as the angle between trochanter-femur and femur-patella segments. The leg movements are shifted to ensure peak acceleration is at 0 ms (vertical dashed line).

## Discussion

Male jumping spiders must execute rapid jumps to escape attacks from females and male competitors during courtship and to avoid predatory attacks. Hence, we focussed on characterising sex-specific differences in the kinematics and choreography of jumps in a peacock spider *M. splendens*. Though we studied only mature adults, the body mass varied between individuals within both males and females, which captured the natural ecological size variation. The body mass variation in males (weight range: 2.0-8.3 mg) and females (weight range: 6.4-13.2 mg) could be the result of slightly different feeding habits, and in addition in females the differing reserves required to invest in eggs. On average, females weighed more than twice as much as males, but both sexes had comparable accelerations and take-off velocities (Fig. 4, Table 1). The smaller sized males were thus able to match the acceleration and take-off velocities of the heavier females, perhaps because smaller body mass means there is less inertia to overcome, and hence less force is required to initiate a jump.

The acceleration of the peacock spider, *M. splendens*, was the fastest among all known jumping spiders; more than twice as fast as the previous highest acceleration in *P. princeps* and more than four times higher than *H. conjunctus*, which had the slowest acceleration among all Salticids (Table 2; [15, 21]). The fast acceleration may be an indication of the predation pressure or the speed of the prey they need to track and capture in natural environments.

We found that both sex and size had a significant effect on all other jump kinematic variables we measured. The lighter males took significantly less time for their take-off compared to the females (Table 1). On average, males and females experienced the equivalent of 13.03 g and 12.5 g respectively during take-off, which is higher than any other known jumping spider (Table 1). In comparison, froghoppers (*Philaenus* sp.) that use stored energy for their jump, experiences 550 g [33]. Since the *g force* experienced by *M. splendens* and other Salticids are several magnitudes lower than animals that rely on stored energy, the semi-hydraulic system that drives jumping in spiders is unlikely to have any elastic or stored energy component. The lighter males also had steeper jump take-off angle compared to the females (Fig. 3B, 3C). We determined take-off angle as the angle of the CoM between two frames: prior to when the abdomen moves and when all legs are off the ground. Although the take-off angle was significantly steeper in males there was a large spread in both sexes for this measure. Surprisingly, the difference in the take-off angle between sexes did not reflect in differences in the maximum height of the jump trajectory (Fig. 3A, 3D, 3E). It could be that the steeper take-off angle and faster take-off durations are adaptations in the lighter males to get away faster from predators. Sexual dimorphism in locomotory strategies is known in spiders and insects. In spiders, small-bodied males outperform heavier females in climbing vertical features or walking faster [31, 32]. In the oriental fruit moth, *Cydia molesta*, females outperformed males in velocity and flight distance [33] and in the butterfly, *Euphydryas phaseton* females flights are sluggish compared to their male counterparts, which is likely due to their greater wing loading which requires a higher wing-beat frequency [34]. In ants, males lead a life on the wing, female workers are exclusively pedestrian, and reproductive females initially lead a life on the wing and subsequently become pedestrian [35]. While sexual dimorphism is prevalent in jumping spiders including *M. splendens*, they have been studied mostly in the context of courtship displays [23, 26] and investigation into their spatial cognition and biomechanics is essential.

### Choreography of leg movements

Peacock spiders are best known for the elaborate courtship display that males carry out to entice the female [25, 26, 36]. As part of this ritual, males of *M. splendens* extend, wave and lower Leg III in front of the female. In contrast to the other legs, Leg III tends to be long and dark with bright white tarsal setae, indicating it is under selection pressure. While Leg III is considered long, actual measurements of the leg have not been reported. Hence, we measured the length and 2D area of leg segments, by dissecting Legs III and IV in one male and female of *M. splendens*. Leg III was 1.23x longer in males compared to the female; Leg IV was 1.04x longer in males compared to the female. In males, Leg III was the longest, with femur being the longest segment. In females, leg IV was the longest, with metatarsus+tarsus section being the longest segment. The 2D area of the femur was largest in the males (Leg III: 386 mm^2^; Leg IV: 348 mm^2^) compared to the female (Leg III: 303 mm^2^; Leg IV: 308 mm^2^). Thus, compared to the female spiders, males invested more in their femurs (being larger in Leg III compared to Leg IV) at least in 2D space.

Legs III and IV played a crucial role in the jumps of both males and females. Well before the start of the jump or displacement of the CoM, in both sexes, spiders raised the first two pairs of legs and extended them in front of their head (Fig. 5; Movie S1). Legs IV left the platform next, and a few milliseconds later was followed by Legs III. In most jumping spiders, similar to *M. splendens*, Legs III are the last to leave the ground. The only exception is *Attulus pubescens*, in which Leg IV leaves the ground last [22]. While the last leg to leave the ground was typically considered as the propulsive leg, Brandt et al., developed an elegant strategy to identify this by calculating the effective leg length (ELL) and angle between leg joints [15]. We determined these two measures relative to the peak acceleration. It was evident that spiders extended and held Legs I and II straight, well before peak acceleration occurred. Leg IV attained the highest ELL and their leg joint was close to a straight angle before the animals attained peak acceleration, with one exception, in all male and female spiders (Fig. 6). This meant the spider continued to accelerate after Leg IV was fully extended and had left the ground. Leg III reached the highest ELL after peak acceleration had occurred. These two findings suggest that Leg III is likely the propulsive leg in both males and females of *M. splendens*, similar to as seen in *H. conjunctus* [15]. Thus, Leg III is under extreme selection since they drive two significant functions: accelerated jumps and courtship behaviours. It remains to be seen, whether males are more vulnerable to predation during courtship since Leg III is extended and waved as part of a visual signal. Equally, it is essential to characterise the 3D volume of the femur and investigate the internal structure and muscles that power the jumps.

### Jump context

Jumping spiders occupy 3D-cluttered environments where they jump under various contexts. Here, we exclusively studied jumps in the context of locomotion that are planned and well executed. Escape or startle jumps provide a strikingly different context compared to locomotory jumps, with the former usually unsteady, non-directed, and tending to have high accelerations. Identifying the kinematics of escape or startle jumps will pinpoint the limits of the system, which locomotory jumps do not capture [6]. Thus, the current known acceleration and velocity in *M. splendens* and *H. conjunctus* are likely less than the maximum kinematics these species can attain [15]. A second and unusual context is multiple jumping or continuous jumping which is a form of endurance locomotion, that jumping spiders and some insects exhibit [37]. Here, we studied only a single jump, and it is unknown how acceleration and jump forces change when animals engage in multiple jumps. A third context is the plane of jumping, i.e., whether an animal carries out horizontal jumps, ascending or descending jumps at different elevations. Here, we studied jumps towards a 4 cm vertical target where animals could land anywhere on the target. Jumping to smaller targets at higher planes will likely require more energy to take-off. Ascending and descending jumps have been described in jumping spiders (*Phidippus regius* and *Phidippus princeps*), but outcomes of these are unclear, since only one individual was studied [16, 21]. In pigeons, potential energy and the power required to fly increases with flight angle [38]. Systematically probing jump kinematics to targets at different elevations would be useful to investigate.

### Conclusions

Salticids exhibit a wide range of body size and occupy different microhabitats. The variation in the jump performance that we described in *M. splendens* is best explained by body mass. We found no difference in take-off velocity and acceleration between sexes, but all other jump kinematics differed between males and females. Jump kinematics have been described in a handful of species, with the smallest spider studied being the males of *M. splendens*. Spiders rely on a combination of hydraulic pressure and muscular action to propel their jump. The jump performance of peacock spiders and other jumping spiders suggest the jump strategy is more similar to the muscle-actuated system and several magnitudes slower compared to the spring-actuated system. A comparative approach to analyse both jump kinematics and hydraulic mechanism that drives the jump is much required. Equally, it will be exciting to analyse the geometry and anatomy of the legs since this can shape the fluid dynamics of leg extension.

## Data and Code Availability

Raw data are available on DRYAD: https://datadryad.org/stash/share/hF4LdhhEM5WhI3xi3oZgxt7KqJUnE-tugm67z0L8vuY.

## Acknowledgements

We acknowledge technical and scientific assistance with micro-CT scanning from Matthew Foley at the Sydney Microscopy & Microanalysis, the University of Sydney.

## Author contributions

AN, DR: Conceptualization; AN, FRE, DR: Methodology; AN, AS, FRE, DR: Investigation; AN, AS, FRE, PJ, DJM, LRO: Analyses; AN, AS, FRE, PJ, DR: Visualization; AN: Funding acquisition, Project administration, and Supervision; AN: Writing – original draft; all authors: Writing – review & editing.

## Figure Legends

Movie S1. Profile videos of jump of the male and female Australian Splendid Peacock spider, *Maratus splendens*. Link to video: https://datadryad.org/stash/share/hF4LdhhEM5WhI3xi3oZgxt7KqJUnE-tugm67z0L8vuY

